# Divergent roles for complement components C3 and C4 in controlling *Klebsiella pneumoniae* gut colonization and systemic dissemination

**DOI:** 10.1101/2025.10.09.681492

**Authors:** Juan D. Valencia-Bacca, Jamie E. Jennings-Gee, Noah A. Nutter, Alexis E. Adams-Sims, Abigail A. Hegarty, Hope L. Didier, M. Ammar Zafar, Karen M. Haas

**Affiliations:** Department of Microbiology and Immunology, Wake Forest University School of Medicine, Winston-Salem, North Carolina 27157, USA; Department of Microbiology and Immunology, Emory University School of Medicine, Atlanta, Georgia 30322, USA

**Author notes:** Co-corresponding authors. Address correspondence to Dr. Karen M. Haas, Wake Forest University School of Medicine, Department of Microbiology and Immunology, 575 North Patterson Avenue, Winston-Salem, NC 27101 and M. Ammar Zafar, Emory University School of Medicine, Department of Microbiology and Immunology, Rollins Research Center, 1510 Clifton Road, Atlanta, GA 30322. **Email:**.

## Abstract

*Klebsiella pneumoniae* is an escalating public health threat driven by the emergence of antibiotic-resistant and hyper-encapsulated strains that spread systemically from the gut. The immune defenses preventing gut colonization and dissemination remain poorly defined. Herein, we uncover distinct and context-dependent roles for complement proteins C3 and C4 in host defense following *K. pneumoniae* infection. Following gut colonization, C3 and C4 levels rise significantly. In addition to inducing alternative pathway-mediated C3b deposition on *K. pneumoniae* grown under gut-relevant conditions, C3 is critical for recruiting myeloid cells to the gut, promoting local opsonophagocytosis, and preventing lethal systemic spread. Depletion of systemic C3 reveals mucosal-derived C3 controls *K. pneumoniae* GI colonization, whereas systemic C3 is essential for limiting fatal dissemination. In contrast, C4 is dispensable for controlling GI colonization, dissemination, and myeloid recruitment under conditions of natural acquisition. However, C4 becomes critical for controlling GI burden and systemic disease following antibiotic-induced dysbiosis and supercolonization with antibiotic-resistant *K. pneumoniae*. Notably, mice deficient in CD21/35—a receptor for cleaved C3 and C4 fragments important for B cell activation and antigen retention—exhibit a defect similar to C4⁻^/^⁻ mice, with significantly increased GI burden under antibiotic-induced supercolonization, suggesting distinct complement-dependent pathways are involved in mucosal protection. Collectively, these findings reveal a dual-layered immune strategy: C3-driven opsonophagocytosis is critical for controlling colonization and dissemination under baseline conditions, while C4 and CD21/35 become indispensable following antibiotic-induced supercolonization. This work advances our understanding of complement-dependent mucosal immune protection and identifies potential targets for preventing gut-to-bloodstream transition of this pathogen.

## Introduction

*Klebsiella pneumoniae* (*Kpn*) is well-recognized as one of the leading causes of antimicrobial-resistant opportunistic infections^1^. *Kpn* can silently colonize the gastrointestinal (GI) tract of healthy individuals without causing clinical manifestations^2–4^. Nonetheless, prolonged presence of virulent strains may contribute to inflammation-driven GI diseases, including cancer^5–8^. Moreover, GI colonization is linked to a heightened risk of subsequent life-threatening extraintestinal infections ^9,10^. In particular, hypervirulent *Kpn* strains which typically exhibit a hypermucoviscous phenotype comprising K1/K2 capsule types, possess an enhanced ability to cross the intestinal mucosal barrier^11,12^. *Kpn* requires a combination of adhesion factors, stress response genes, and genetic regulatory networks to translocate from the GI tract and establish extraintestinal infections^13^ –a process that may entail transcellular invasion without disrupting tight junctions or epithelial integrity^14^. Identifying host defense mechanisms that protect against *Kpn* colonization, translocation, and dissemination are critical for devising strategies to protect against *Kpn* infections threatening human health.

Host defense against bacterial colonization in the gut relies on multiple factors, including the microbiota, mechanical barriers, and innate and adaptive immune responses^15^. Among these, innate immunity plays a pivotal role as the first line of defense, detecting microbial invaders through pattern recognition receptors (PRRs) and rapidly initiating phagocytic and inflammatory responses to restrict the expansion and translocation of harmful, non-commensal microorganisms^16^. Despite this, the role of the complement system—a key element of innate immunity, in protecting against intestinal infections remains unclear. While the liver is the main source of complement proteins^17,18^, non-hepatic sources, such as immune cells, intestinal epithelial cells, and colonic stromal cells, also synthesize these components^19–21^. The function of the complement system in the gut has been largely overlooked, partly due to the limited detection of complement components in the healthy GI lumen^20^. This perception is further reinforced by the prominent role of secretory IgA (SIgA) in gut immunity, which, despite its effectiveness in promoting non-inflammatory clearance of pathogens and maintaining mucosal homeostasis, has a limited capacity to activate complement^22^. As a result, few studies have specifically investigated the role of complement in gut colonization and subsequent systemic dissemination, leaving significant gaps in our understanding of its contribution to early-stage host defense. However, a recent study by Wu et al. showing a critical role for the C3-dependent alternative pathway in providing protection against *Citrobacter rodentium* GI infection in mice has ignited interest in this area^21^.

Reports on the susceptibility of *Kpn* strains to complement-mediated killing and clearance are variable. While serum-sensitive strains activate both the classical and alternative complement pathways *in vitro*, leading to C3b deposition and subsequent lysis via membrane attack complex (MAC) formation^23^, serum-resistant strains evade MAC-mediated lysis primarily due to a thick capsule and/or extended O-antigen polysaccharide chains in lipopolysaccharide (LPS), which can redirect C3b deposition away from the bacterial membrane ^24,25^. Notably, in clinical isolates lacking the LPS O antigen, the capsular type has been shown to be a key determinant of complement resistance^26^. Interestingly, a recent study demonstrated K23 capsule-expressing *Kpn* were cleared by a C3-independent mechanism involving liver-resident Kupffer cells^27^. Mouse models of *Kpn* lung infection investigating C3 effects using the virulent K2 capsule-type KPPR1 (ATCC 43816) *Kpn* strain have shown contradictory roles, with one study showing increased splenic dissemination in C3^-/-^ mice^28^, whereas another showed no differences in survival relative to wildtype mice^29^. Thus, the role of complement in controlling *Kpn* infection remains unclear. In particular, the role of complement in controlling *Kpn* gut colonization and subsequent systemic dissemination is completely unknown. Herein, we demonstrate context dependent roles for C3 and C4 in controlling *Kpn* GI-associated pathogenesis.

## Results

### Complement component C3, but not C4, is required to limit *K. pneumoniae (Kpn)* burden in the gut following natural acquisition

Oral inoculation of *Kpn* strain KPPR1S (ATCC 43816, ST493, K2 serotype) leads to robust GI colonization in wild-type (WT) mice with no noticeable clinical effects^30^. However, it is possible *Kpn* colonization induces sub-clinical inflammation. We therefore measured C3 and C4 complement protein levels in fecal samples from naive C57BL/6 WT mice and at 24, 48, 72 and 96 hours post-oral *Kpn* inoculation (Fig. 1A-B), along with fecal shedding (Fig. 1C). Consistent with previous studies, fecal samples from naive mice exhibited low basal or undetectable levels of C3 and C4^21^. Conversely, at 72- and 96-hours post-inoculation, C3 levels significantly increased (>5-fold; Fig. 1A), and by 96 hours, C4 levels were also significantly increased (Fig. 1B) suggesting a mucosal immune response. We therefore determined whether C3 and/or C4 influence bacterial burden in the GI tract by evaluating *Kpn* GI colonization in WT, C3⁻^/^⁻, and C4⁻^/^⁻ mice. At 24 hours post-inoculation, C3⁻^/^⁻ mice, but not C4⁻^/^⁻ mice, exhibited a significantly higher bacterial burden (1000-fold increased) in feces relative to WT mice (Fig. 2A). At 24 hours post challenge, recovered CFU was also significantly increased (1000-fold) at all sites sampled along the GI tract (ileum, cecum, and colon) of C3⁻^/^⁻ mice, whereas C4⁻^/^⁻ burden was similar to WT mice (Fig. 2B). This striking phenotype persisted up to 96 hours post-challenge (Fig. 2C). Thus, C3, but not C4, is critical for limiting *Kpn* intestinal burden following oral challenge.

**Figure 1.**
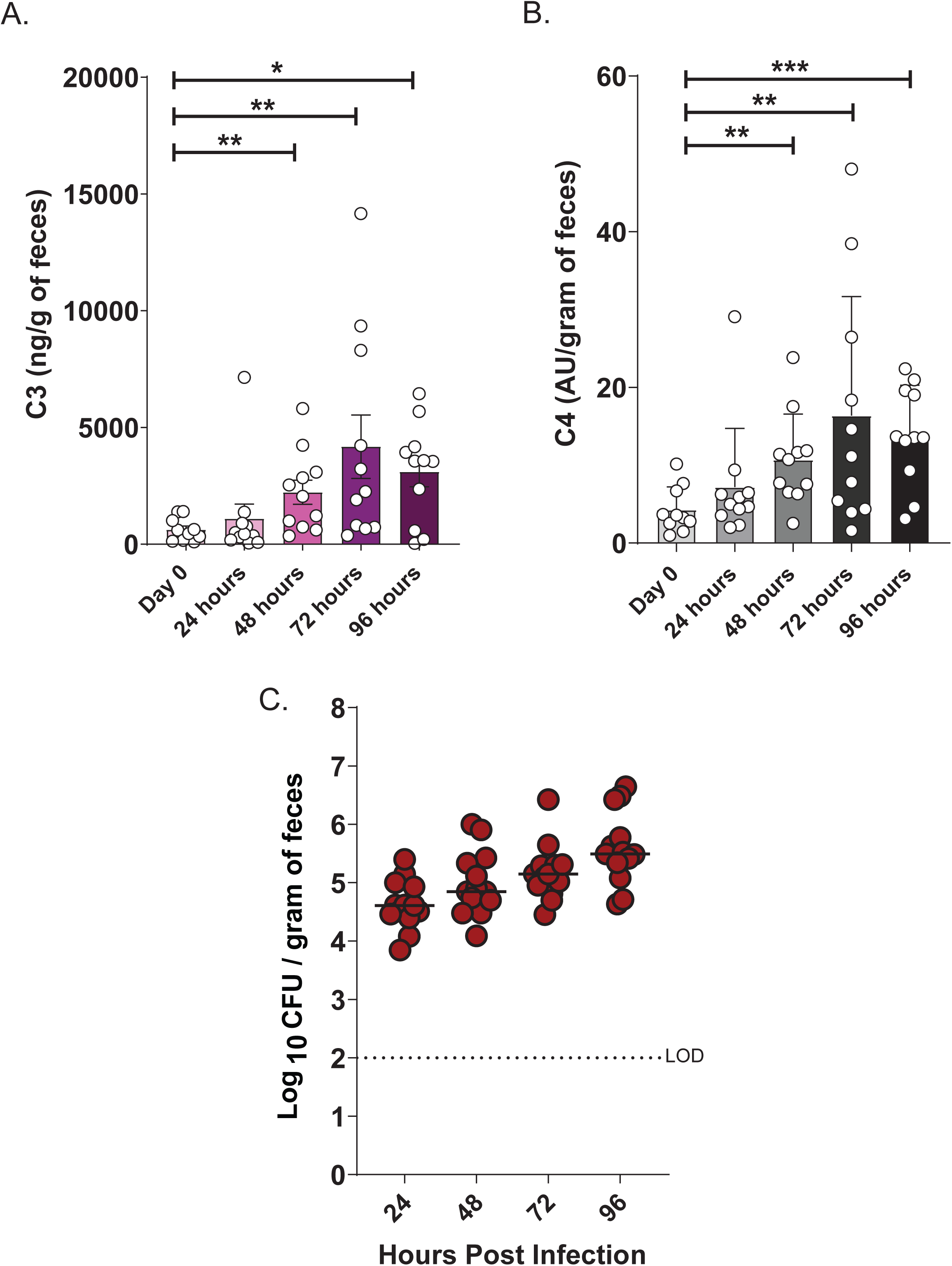
Fecal C3 and C4 levels following gastrointestinal colonization with *Kpn*. (A–B) C3 (A) and C4 (B) levels measured by ELISA in fecal samples from C57BL/6J wild-type (WT) mice before infection, and at 24, 48, 72 and 96 hours following oral inoculation with *Kpn* strain KPPR1S (∼10⁶ CFU). C) Corresponding fecal shedding in mice colonized with KPPR1S. Each symbol represents an individual mouse (n =13 per group). Statistical significance was assessed by one-way ANOVA with Bonferroni post hoc test (A-B). *\*P* < 0.05, *\*\*P* < 0.01, *\*\*\*P* < 0.001, and *\*\*\*\*P* < 0.0001.

**Figure 2.**
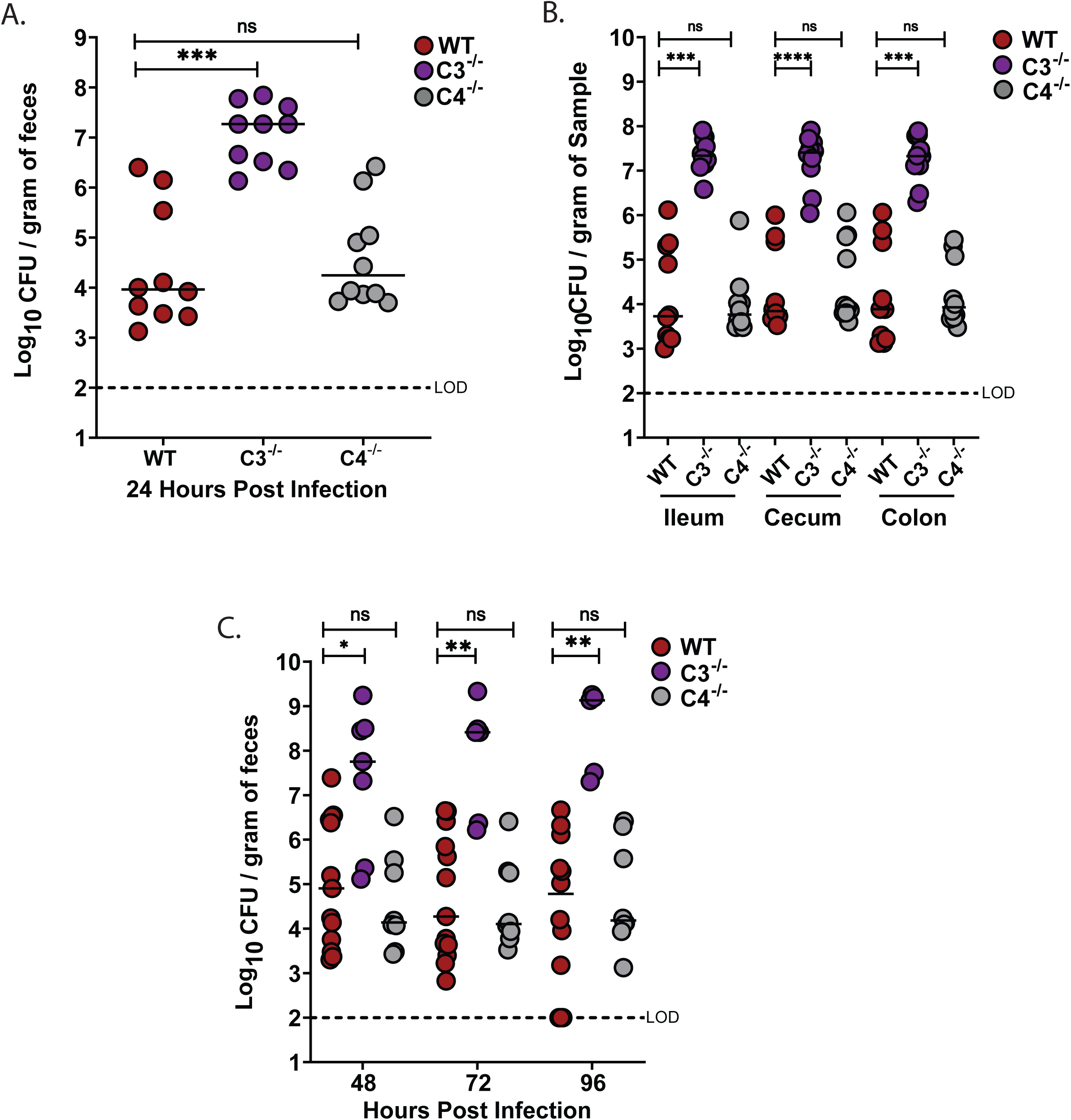
C3, but not C4, controls *Kpn* gut burden, systemic dissemination, and survival following oral inoculation. (A) Fecal shedding in mice colonized with KPPR1S 24 hours post-infection. (B) Bacterial densities in the ileum, cecum, and colon at 24 hours. (C) Fecal shedding at 48, 72, and 96 hours. Each circle represents an individual WT (red), C3^⁻/⁻^ (purple), or C4^⁻/⁻^ (gray) mouse. Dotted lines indicate the limit of detection (L.O.D.). Similar results were obtained in two independent experiments with 4-5 mice per genotype. Pooled data are shown. Statistical significance was assessed by Kruskal-Wallis with Dunn’s post-hoc test (A–B); n ≥ 9, (C). n ≥ 7. *\*P* < 0.05, *\*\*P* < 0.01, *\*\*\*P* < 0.001, and *\*\*\*\*P* < 0.0001.

### Complement component C3, but not C4, is required to limit *Kpn* extraintestinal dissemination and death

Molecular epidemiological studies suggest *Kpn* gut colonization is a marker for subsequent invasive disease^9,10^. We therefore assessed the presence of *Kpn* in the liver, spleen and blood of WT, C3⁻^/^⁻, and C4⁻^/^⁻ mice. At 24 hours post-inoculation, approximately half of the WT and C4⁻^/^⁻ mice harbor *Kpn* in the liver (Fig. 3A), and to a lesser extent in the spleen and blood. However, all C3⁻^/^⁻ mice were colonized with *Kpn* in the liver, spleen, and blood, with bacterial densities ∼1000-fold over that found in WT and C4⁻^/^⁻ mice. Consequently, the majority of C3⁻^/^⁻ mice succumbed to infection, in contrast to WT and C4⁻^/^⁻ mice (Fig. 3B). Thus, in addition to controlling *Kpn* GI colonization and extraintestinal burden following oral *Kpn* colonization, C3, but not C4, is required for survival after dissemination.

**Figure 3.**
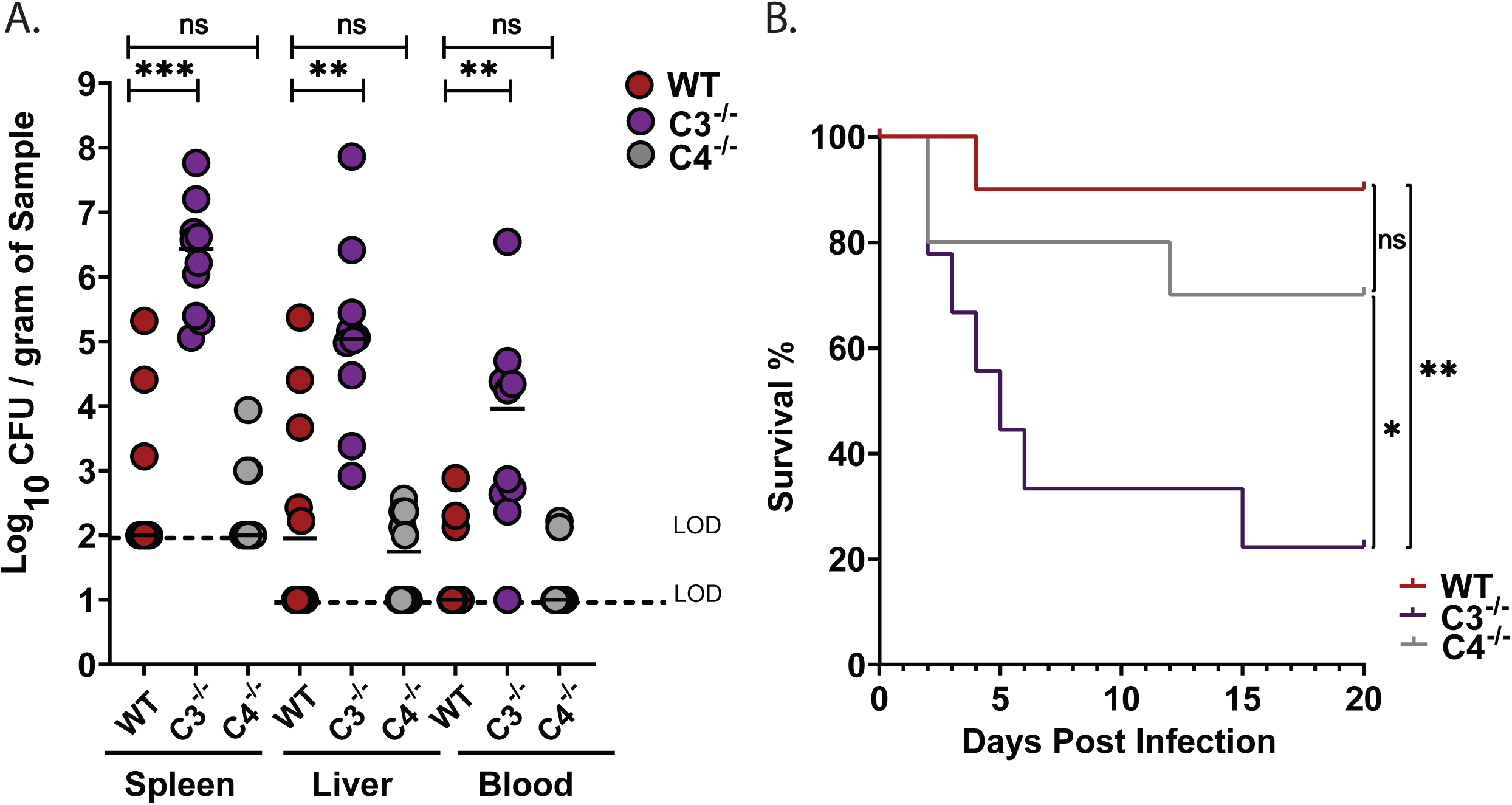
Systemic dissemination and survival following oral *Kpn* infection. (A) Bacterial burden in spleen, liver, and blood at 24 hours. (B) Kaplan-Meier survival curve indicating survival following oral KPPR1S inoculation. Each circle represents an individual WT (red), C3^⁻/⁻^ (purple), or C4^⁻/⁻^ (gray) mouse. Dotted lines indicate the limit of detection (L.O.D.). Similar results were obtained in two independent experiments with 4-5 mice per genotype. Pooled data are shown. Statistical significance was assessed by Kruskal-Wallis with Dunn’s post-hoc test and the Log-rank (Mantel-Cox) tests (B). (A-C); n ≥ 9. *\*P* < 0.05, *\*\*P* < 0.01, *\*\*\*P* < 0.001, and *\*\*\*\*P* < 0.0001.

### *Kpn* is resistant to complement-mediated lysis under *in vitro* gut-mimicking growth conditions but remains sensitive to C3b deposition in a C4-independent manner

The inability of C3⁻^/^⁻ mice to control *Kpn* gut burden could be due to defects in complement-mediated opsonophagocytosis and/or the ability of the complement system to directly kill gram-negative bacteria through MAC formation. To investigate potential mechanisms by which C3 contributes to reduced gut burden, we assessed complement-mediated effects on *Kpn* grown in either gut mimetic media (cecal filtrate) or lysogeny broth (LB). Bacteria were incubated with 10% rabbit complement for 30 or 60 minutes, and survival was determined. As shown in Figure 4A-B, KPPR1S remained completely resistant to complement-mediated killing regardless of whether it was cultured in LB or cecal filtrate, with no significant reduction in CFU relative to 10% heat-inactivated complement or PBS alone, consistent with a previous report^25^. In contrast, an isogenic capsule-deficient strain (*ΔwcaJ*) was highly susceptible to complement-mediated killing, with 10% rabbit complement reducing recovered CFU 1000-fold (Supplementary Fig. 1A-B). Heat-inactivated complement did not affect bacterial survival relative to PBS, confirming that the killing was specifically mediated by active complement components. Although KPPR1S was not killed by 10% active rabbit complement, it induced significant C3b deposition on KPPR1S grown in either LB or cecal filtrate (Fig. 4C and 4D). In contrast, no C3b deposition was observed in the presence of heat-inactivated complement. Pretreatment of rabbit complement with EGTA, which chelates calcium ions and selectively inhibits the classical and lectin pathways, modestly reduced C3b deposition when *Kpn* was grown in LB (Fig. 4E) but had no detectable effect on C3b deposition for bacteria grown in cecal filtrate (Fig. 4F). These findings suggest that the alternative pathway may be a major driver of C3b deposition under gut-mimicking conditions, and may in part, explain the lack of effect of C4-deficiency on *Kpn* colonization levels.

**Figure 4.**
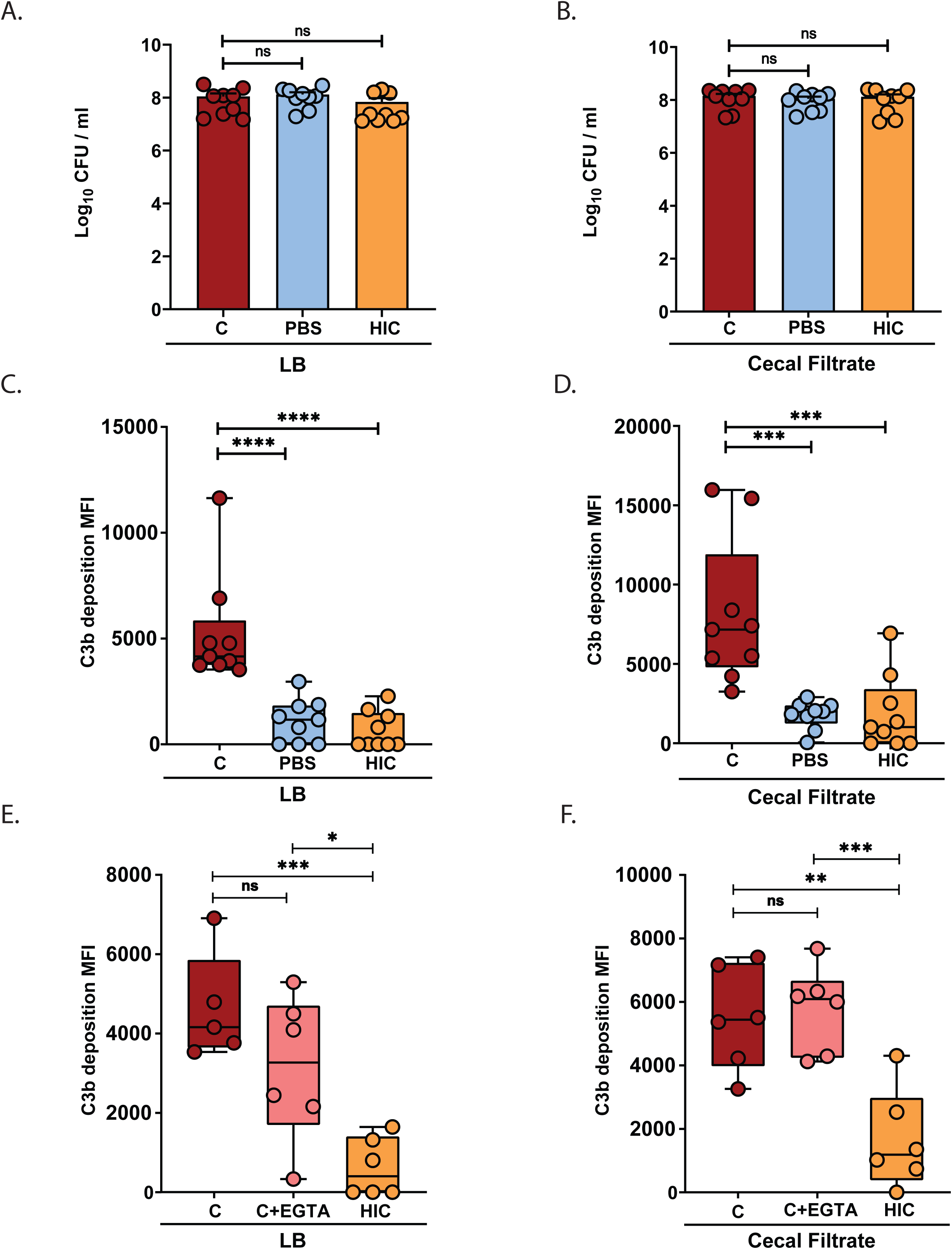
*Kpn* induces C3-dependent, C4-independent, C3b deposition and myeloid cell recruitment to the gut. (A–B) *Kpn* KPPR1S was incubated for 1 hour at 37 °C in LB (A) or cecal filtrate (B) in presence of 10% rabbit complement, heat-inactivated complement (HIC), or PBS. Recovered CFU was assessed, and significance was determined using the Kruskal-Wallis test with Dunn’s post-hoc test. (C-E) C3b deposition on KPPR1S grown in LB or cecal filtrate was analyzed by flow cytometry after 1-hour incubation at 37 °C. (C-F) Flow cytometry quantification of C3b deposition on KPPR1S in LB (C, E) or cecal filtrate (D, F) with 10% rabbit complement, EGTA-treated complement, or HIC. A-D, Pooled results from 3 independent experiments. E-F, Pooled results from 2 to 3 independent experiments. Statistical analysis: one-way ANOVA with Bonferroni post hoc test (n≥ 5 per group). *\*P* < 0.05, *\*\*P* < 0.01, *\*\*\*P* < 0.001, and *\*\*\*\*P* < 0.0001.

### *Kpn* GI colonization induces C3-dependent myeloid cell recruitment into the colon

In addition to supporting C3b-mediated phagocytosis, C3 activation can support leukocyte recruitment into sites of inflammation^31^. We assessed leukocyte numbers in the colon by flow cytometry 48 hours after oral inoculation and found a significant increase in CD45⁺ cells in infected WT and C4⁻^/^⁻ mice (Fig. 5A), which was concomitant with a 5-fold increase in CD45⁺GR-1⁺SSC^hi^ (Fig. 5B), indicative of myeloid cell recruitment^32^. Interestingly, baseline levels of CD45^+^ and GR1^+^ cell numbers were lower in C3⁻^/^⁻ than WT mice, and in contrast to WT and C4⁻^/^⁻ mice, we did not observe a significant increase in these cells following oral inoculation in C3⁻^/^⁻ mice. Thus, C3, but not C4 is required for *Kpn* opsonophagocytosis by inducing C3b deposition as well as by supporting phagocyte recruitment to the intestine following infection.

**Figure 5.**
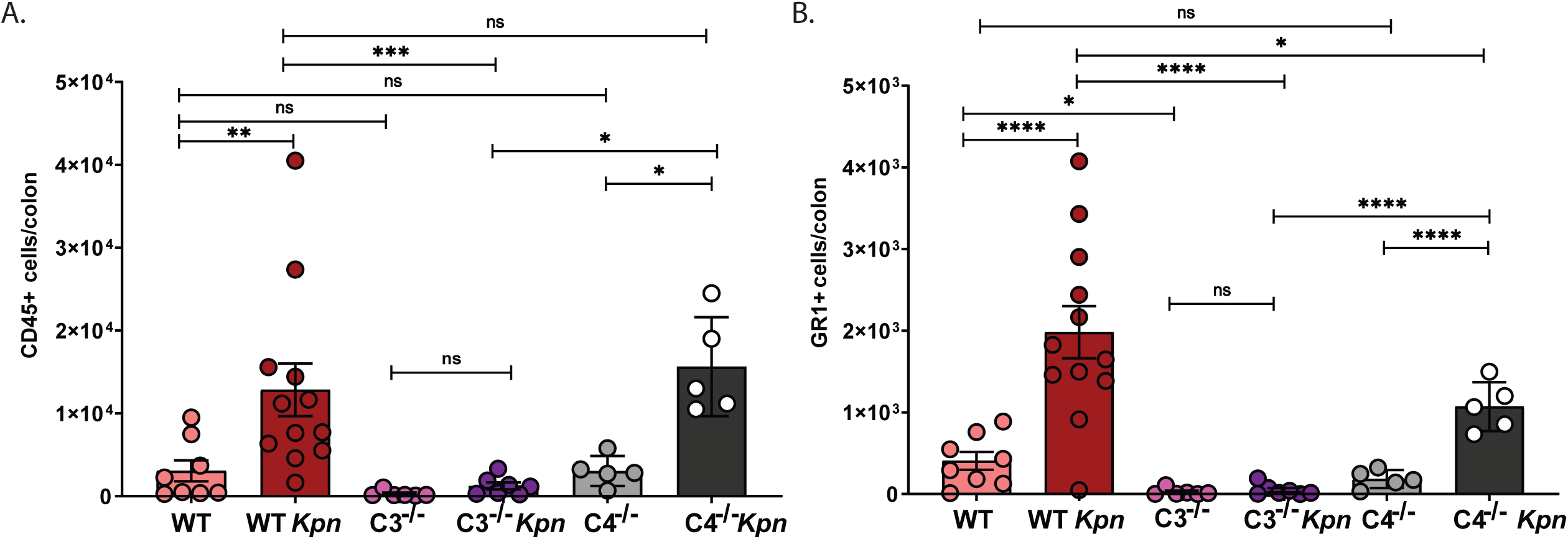
Colonic leukocyte and myeloid cell infiltration following oral *Kpn* inoculation. (A-B) Flow cytometry detection of CD45^+^ leukocyte (A) and GR1^+^ myeloid (B) cells in the colons of naïve control mice and at 48 hours post-oral inoculation with KPPR1S. Pooled results from 2-3 independent experiments, with each circle indicating results from an individual mouse. Statistical outliers associated with sample processing issues and poor colonization were removed in GraphPad (Rout, Q=1%) in G-H. Statistical analysis was conducted using two-way ANOVA (n≥ 5 mice per group). *\*P* < 0.05, *\*\*P* < 0.01, *\*\*\*P* < 0.001, and *\*\*\*\*P* < 0.0001.

### Control of *Kpn* GI burden and dissemination in the context of antibiotic-induced supercolonization requires both C3 and C4

In clinical settings, prolonged antibiotic therapy is a significant risk factor for developing severe and invasive *Kpn* infections^33^. Antibiotic use reduces microbial diversity in the GI tract and facilitates high level colonization of antibiotic-resistant *Kpn*^34^. To investigate the role of complement in *Kpn* GI colonization and dissemination in the context of antibiotic treatment, we employed a broad-spectrum antibiotic treatment model using streptomycin, delivering the bacterial load via orogastric gavage. Antibiotic treatment results in supercolonization of the streptomycin-resistant KPPR1S strain, with colonization levels increased 1000-fold over that found with oral inoculation in the absence of antibiotic treatment^30^. Similar to our results obtained with oral feeding, with antibiotic treatment C3⁻^/^⁻ mice displayed 100-to 1000-fold greater *Kpn* CFU in fecal shedding, cecum, and colon relative to WT mice (Fig. 6A-C). In contrast to our results with natural colonization whereby C4 deficiency had no observable effect on *Kpn* gut burden, with antibiotic treatment, C4⁻^/^⁻ mice had significantly increased fecal shedding, cecum, and colon CFUs relative to WT mice, and at levels that were comparable to C3⁻^/^⁻ mice (Fig. 6A-C).

**Figure 6.**
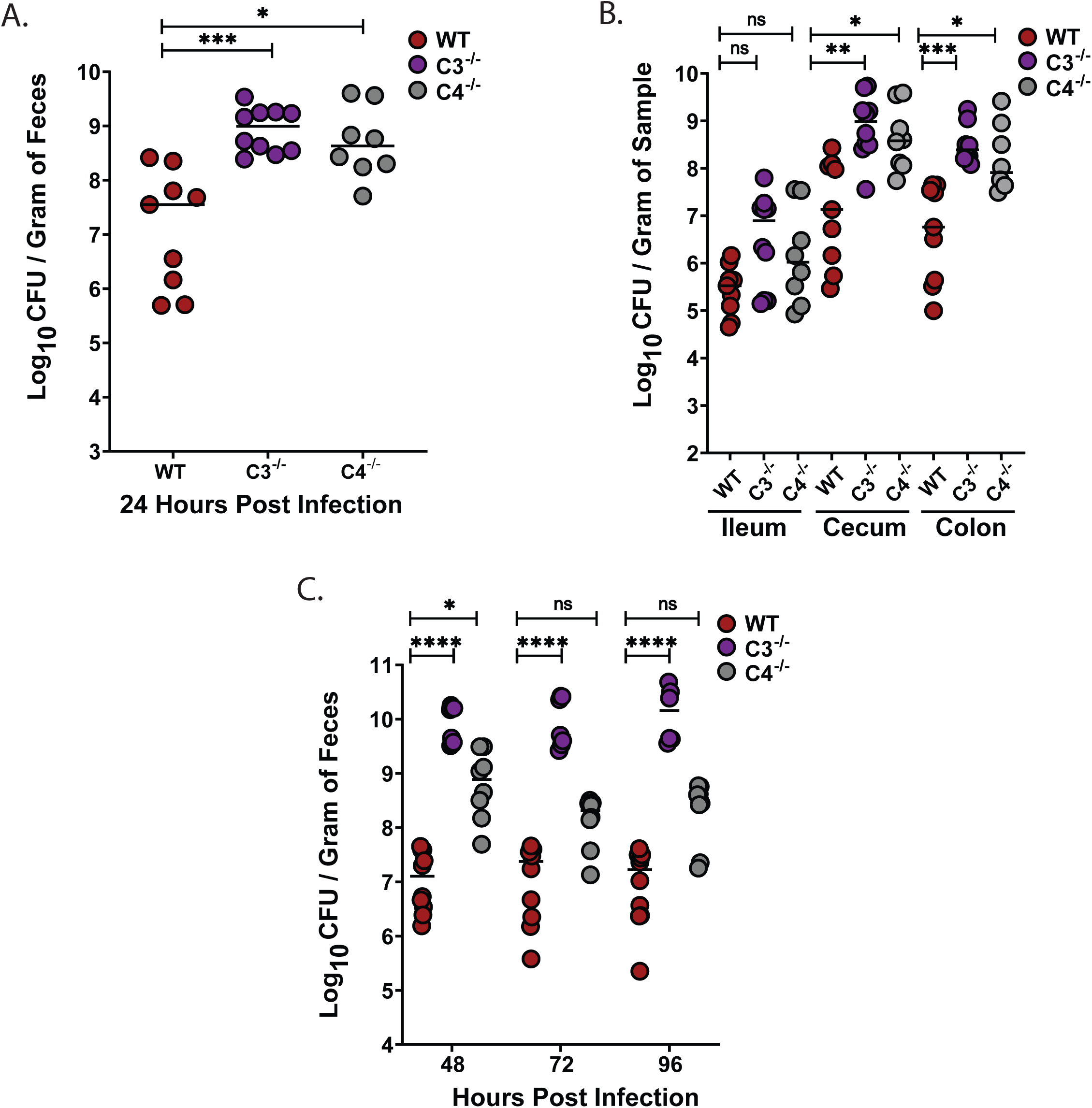
C3 and C4 are both required to control *Kpn* GI burden in the context of antibiotic-induced supercolonization. (A) Fecal shedding of KPPR1S at 24 hours post-gavage in streptomycin-pretreated mice (500 µg/mL in drinking water). (B) KPPR1S bacterial loads in the ileum, cecum, and colon at 24 hours post-gavage. (C) Fecal shedding of KPPR1S at 48, 72, and 96 hours post-gavage. Each point in panels A–C, represents an individual mouse. Dotted lines indicate the limit of detection (L.O.D.). All bacterial strains were grown overnight in LB, and mice were inoculated with ∼10⁷ CFU. A-C, Pooled results from 2-3 independent experiments, with each circle indicating results from an individual mouse. Statistical significance was determined using the Kruskal–Wallis test with Dunn’s post hoc test (A–C); n ≥ 9/group). *\*P* < 0.05, *\*\*P* < 0.01, *\*\*\*P* < 0.001, and *\*\*\*\*P* < 0.0001.

Relative to WT mice, C3⁻^/^⁻ and C4⁻^/^⁻ mice had higher bacterial loads in extra-intestinal sites (spleen, liver, and blood) at 24 hours post infection (Fig. 7A). Consistent with this, survival was significantly reduced in highly colonized C3⁻^/^⁻ and C4⁻^/^⁻ mice relative to WT mice (Fig. 7B). Nonetheless, survival analysis in the antibiotic treatment model highlights the pivotal role of C3 in controlling systemic dissemination, with survival rates of less than 10% in C3⁻^/^⁻ mice, compared to ∼75% in C4⁻^/^⁻ and 100% in WT mice (Fig. 7B).

**Figure 7.**
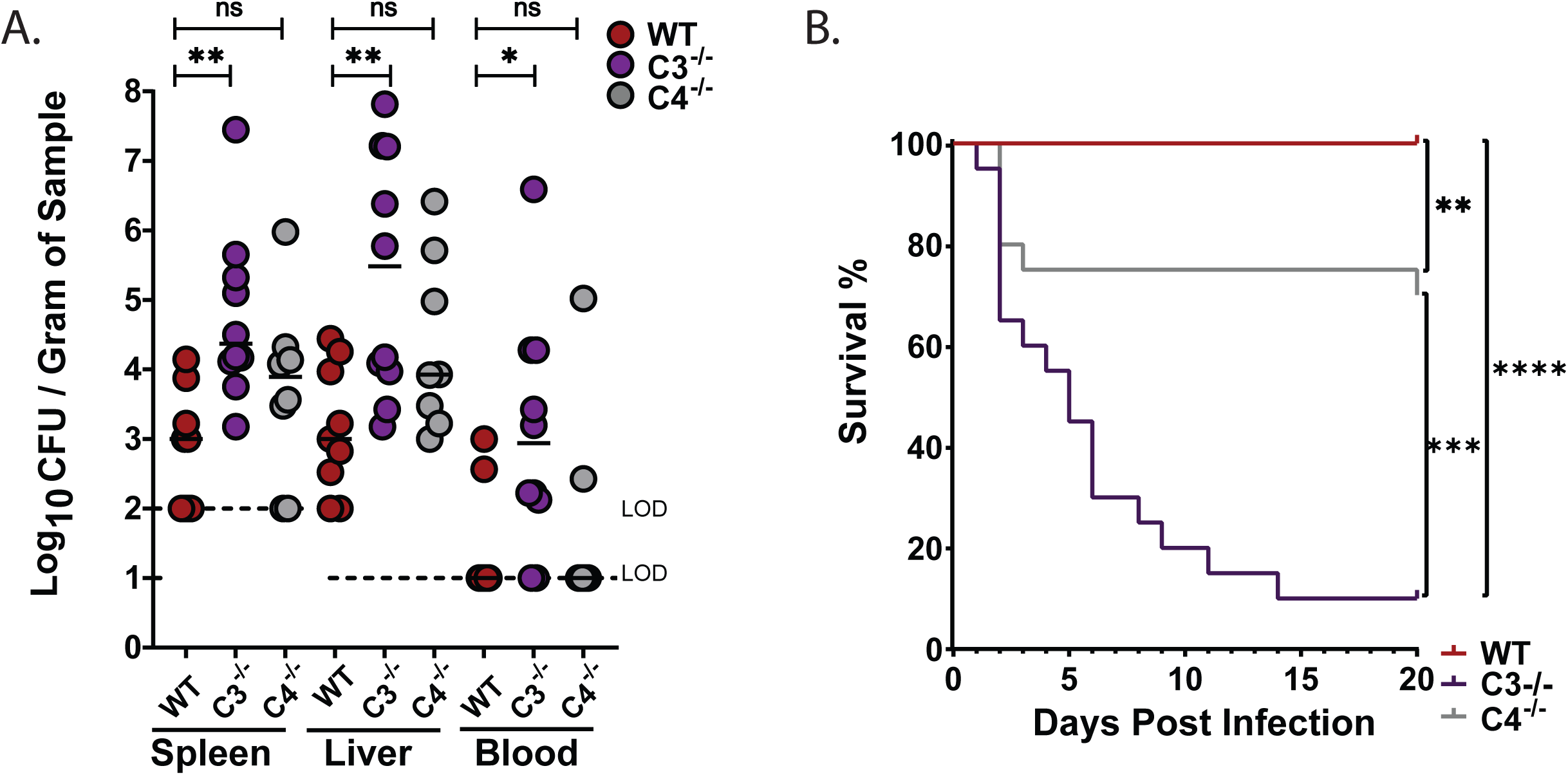
C3 and C4 are essential to prevent dissemination and mortality during antibiotic-driven supercolonization. (A) Systemic dissemination assessed by bacterial burden in spleen, liver, and blood at 24 hours post-gavage. (B) Kaplan–Meier survival curves of WT (red), C3^-/-^ (purple), and C4^-/-^ (gray) mice following KPPR1S oral gavage. (Statistical significance was determined using the Kruskal–Wallis test with Dunn’s post hoc test (A) n ≥ 9/group, and the Log-rank (Mantel-Cox) tests (B), n ≥ 20. *\*P* < 0.05, *\*\*P* < 0.01, *\*\*\*P* < 0.001, and *\*\*\*\*P* < 0.0001.

### Depletion of circulating C3 in WT mice has no effect on *Kpn* GI colonization but leads to fatal dissemination

To distinguish the roles of systemic versus locally produced C3 in protection against invasive disease, we used cobra venom factor (CVF) to deplete circulating C3 while preserving GI-derived C3^21^. Depletion of systemic C3 activity in WT mice had no effect on *Kpn* burden in the gut (Fig. 8A). However, a significant fraction of CVF-treated WT mice succumbed to bacterial dissemination (Fig. 8B), indicating that circulating complement is critical for controlling systemic infection but is dispensable for regulating bacterial colonization in the GI tract.

**Figure 8.**
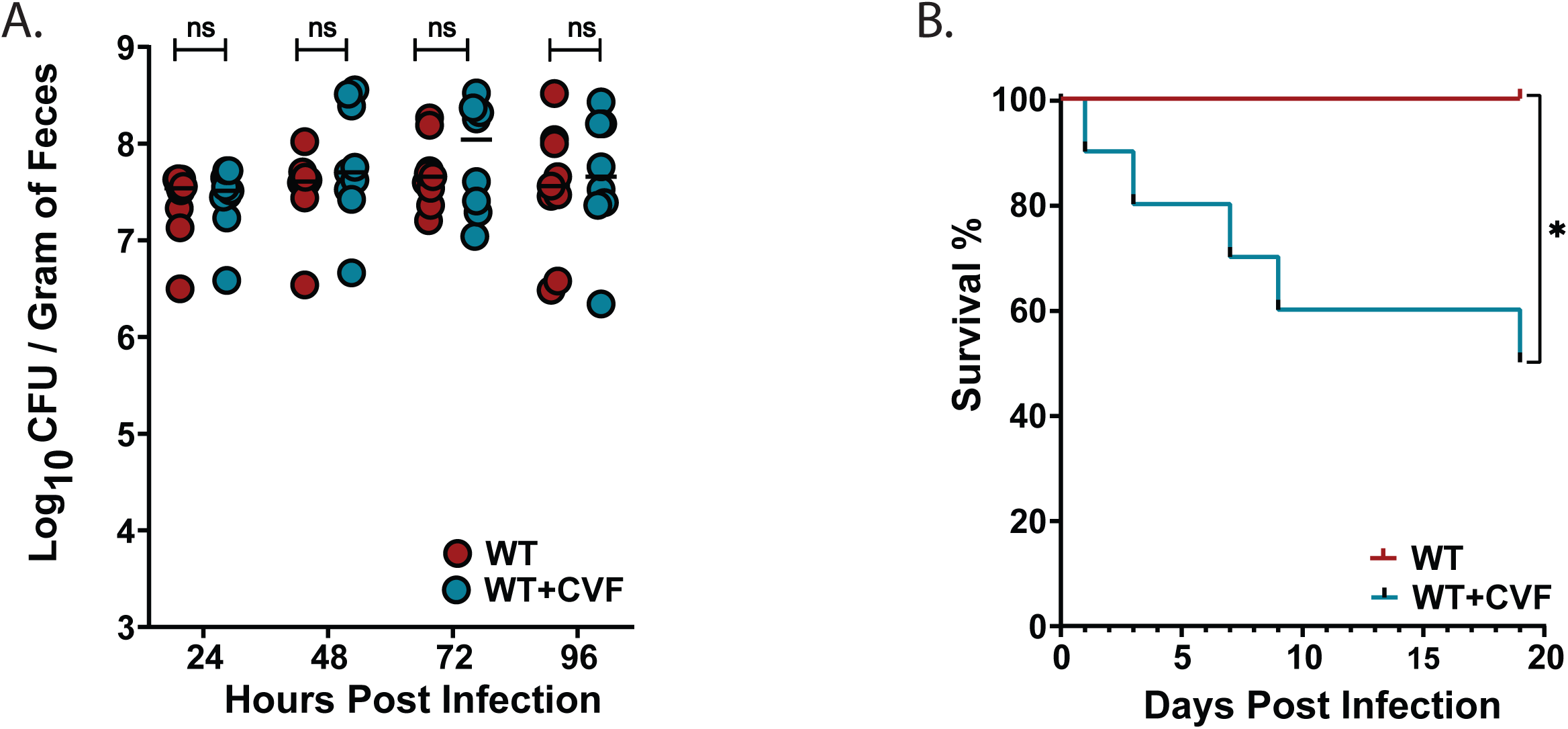
Depletion of circulating C3 in WT mice increases fatal dissemination but has no effect on *Kpn* GI colonization levels. (A) Fecal shedding in WT (red) and CVF-treated WT mice (blue) at 24, 48, 72, and 96 hours post-KPPR1S gavage inoculation. (B) Kaplan–Meier survival of WT and CVF-treated mice following oral KPPR1S gavage. Statistical significance was determined using Mann–Whitney U test (A) for bacterial burden; n ≥ 9/group), and the Log-rank (Mantel-Cox) tests (B) for survival (G: n ≥ 9). *\*P* < 0.05, *\*\*P* < 0.01, *\*\*\*P* < 0.001, and *\*\*\*\*P* < 0.0001.

### C3 and C4 are required for control of antibiotic-induced supercolonization of classical *Kpn* strain MKP103

Due to genetic heterogeneity of *Kpn* isolates^35^, we investigated whether GI colonization with the classical multi-drug resistant *Kpn* strain MKP103—a derivative of the outbreak strain KPNIH1 (ST258)^36^, is similarly regulated by C3 or C4 in the context of ampicillin treatment. While no differences in colonization were observed at 24 hours, MKP103 GI burden was significantly higher in both C3⁻^/^⁻ and C4⁻^/^⁻ mice at 48 hours, and by 96 hours was >100-fold increased over WT mice (Supplementary Fig. 2A). However, all MKP103-colonized mice survived the infection (Supplementary Fig. 2B), consistent with reports of its limited virulence in mice^37^. Thus, in contrast to its limited role in controlling *Kpn* colonization and extra-intestinal dissemination following natural acquisition, in the context of antibiotic-induced *Kpn* supercolonization, C4 is required along with C3 to limit bacterial burden in the GI tract for both hypervirulent and classical strains.

### The C3/C4 fragment complement receptor, CD21/CD35, is required for controlling antibiotic-induced *Kpn* GI supercolonization

The significantly increased *Kpn* GI burden in C4⁻^/^⁻ mice despite finding a limited role for C4 in C3b-mediated deposition or myeloid cell recruitment suggested distinct immunologic defects may exist with C4 deficiency that result in unchecked GI burden under antibiotic conditions. C4 is well known to promote antibody production^38^, although its role in supporting mucosal antibody production is not well-investigated. We therefore investigated *Kpn* colonization in mice lacking complement receptors CD21 and CD35 (complement receptors 1 and 2), which recognize cleaved products of C3 and C4 and support B cell activation, germinal center formation, and long-term humoral immunity, particularly through their role in enhancing B cell receptor signaling and antigen retention on follicular dendritic cells^39–43^. Notably, we found CD21/35-deficient mice have significantly increased *Kpn* GI burden under antibiotic-induced KPPR1 supercolonization, similar to C4^-/-^ mice (Fig. 9A), although survival was comparable to WT mice (Fig. 9B). Together, these findings suggest that the C4-CD21/35 axis plays a previously underappreciated role in shaping mucosal immunity which renders protection against the high levels of *Kpn* which occur under conditions of dysbiosis, where distinct complement-dependent mechanisms in addition to opsonophagocytosis may become essential for maintaining barrier integrity and limiting pathogen overgrowth.

**Figure 9.**
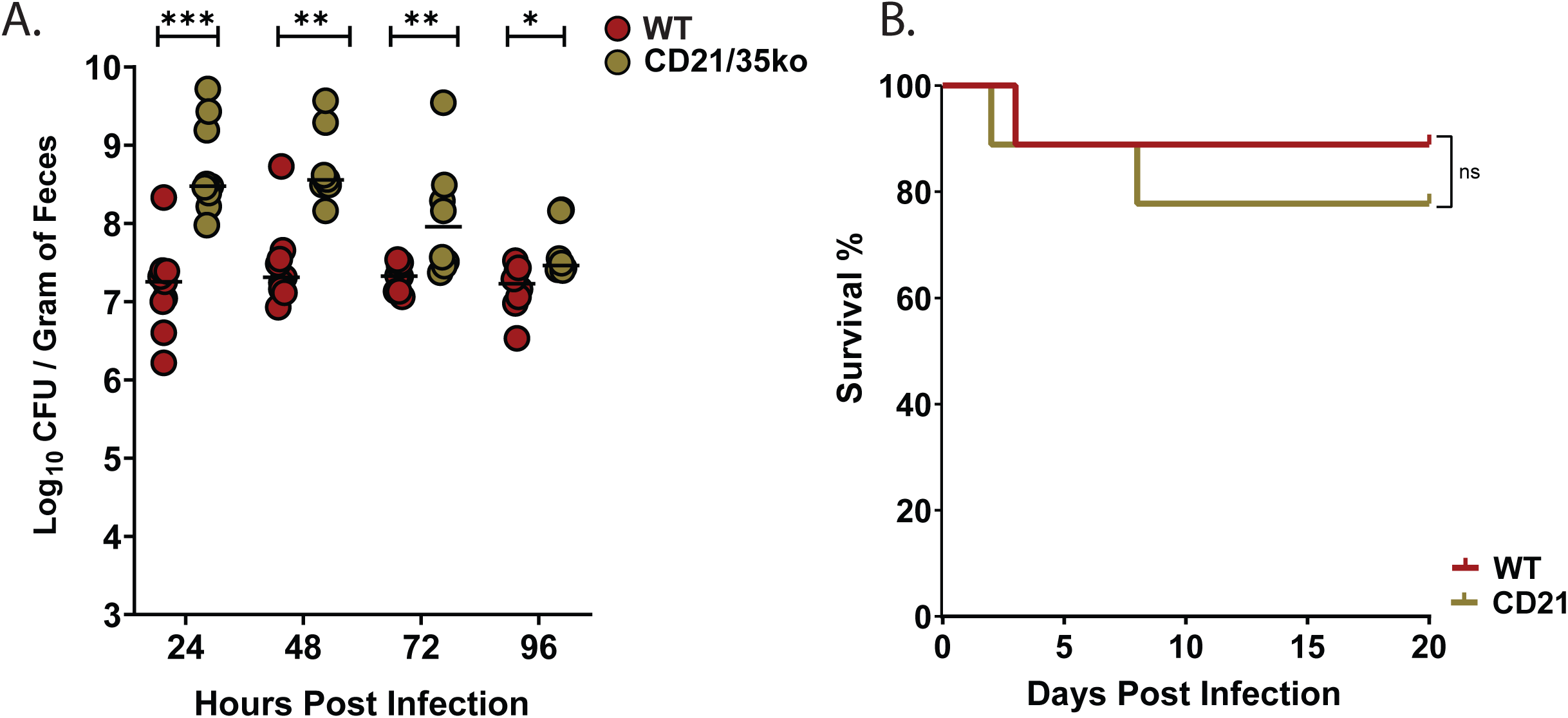
CD21/35 is required to control *Kpn* GI burden in the context of antibiotic-induced supercolonization. A) Fecal shedding of KPPR1S at 24,48,72 and 96 hours post-gavage in streptomycin-pretreated (500 µg/mL in drinking water), WT mice (red circles) and CD21/35^-/-^ mice (golden circles). (B) Kaplan–Meier survival curves of WT and CD21/35^-/-^ mice following KPPR1S oral gavage. Statistical significance was determined using the Kruskal–Wallis test with Dunn’s post hoc test (A) n ≥ 9/group, and the Log-rank (Mantel-Cox) tests (B), n ≥ 20. *\*P* < 0.05, *\*\*P* < 0.01, *\*\*\*P* < 0.001, and *\*\*\*\*P* < 0.0001.

## Discussion

The complement system, an evolutionary ancient component of the innate immune response plays a fundamental role in protecting the host against microbes that breach barriers and enter the bloodstream^44,45^. However, emerging evidence points to a role for complement production at these barrier surfaces, including the gastrointestinal and respiratory tract^17,46–52^. The data presented herein reveals several insightful findings regarding the role of complement in *Kpn* pathogenesis. First, we reveal an unexpected role for locally-produced C3 in its ability to limit *Kpn* colonization and expansion within the gastrointestinal tract. Second, we demonstrate that despite its ability to prevent lysis, the K2 capsule produced under GI growth conditions allows for C3-deposition to occur via the alternative pathway. Third, besides providing protection against systemic disease, C3 is critical for controlling GI burden through phagocyte recruitment in conjunction with its capacity for opsonization. Finally, we find C4—central to the classical and lectin pathways—is dispensable for controlling GI burden and dissemination at low colonizing burdens encountered in the absence of selective antibiotics; however, under conditions of antibiotic-induced *Kpn* bloom in the gut, C4 becomes critical for controlling GI burden, potentially through a pathway requiring the C3b/C3d/C4b receptor, CD21/CD35. As *Kpn* GI burden demonstrates a strong correlation with the degree of translocation (Supplemental Fig. 3A) and dissemination, accordingly, complement-mediated control of GI burden, whether it be C3-dependent/C4-independent in the context of natural colonization or C3- and C4-dependent in the context of antibiotic treatment, is a critical first line of defense against *Kpn* systemic disease.

The finding that C3 restricts *Kpn* gut burden is consistent with recent findings reported by Wu et al. for *Citrobacter rodentium* infection^21^. While their results suggested infection-induced neutrophil recruitment induces mucosal C3 levels and is associated with alternative pathway-mediated phagocytosis, our data directly demonstrates C3 as being critical for initial recruitment of phagocytic cells to the infection site and thereby suggests a broader role for C3 in defense of the gastrointestinal tract in the context of *Kpn* infection. While we did not identify the source of mucosal C3, the initial low levels likely produced by colonic stromal cells in non-infected mice^21^, may be critical for myeloid cell recruitment, as is evidenced by the reduced number of Gr1^+^ cells isolated from colons of non-infected C3^-/-^ mice. The extent to which local C3a-responsive cells, such as mast cells and/or epithelial cells, as well as *Kpn* LPS-induced upregulation of C3aR on these and other cell types^53^, is involved in eliciting signals that may be required for myeloid cell recruitment requires further study.

The critical role for C3 relative to C4 in regulating dissemination stems partly from its role in controlling GI burden. C3b deposition by the alternative pathway is likely critical both in the GI tract and in the circulation, as supported by our CVF depletion results, and the increased bacterial burden and susceptibility of C3⁻^/^⁻ mice to systemic challenge relative to C4⁻^/^⁻ mice. Importantly, our CVF depletion results suggest a critical role for circulating C3 in limiting systemic *Kpn* dissemination, distinct from its role in controlling GI burden. Expression of C3b receptors (CrIg, CD11b/CD18) by Kupffer and other phagocytic cells provides a likely path toward *Kpn* clearance^54,55^,. Although circulating *Kpn*-reactive natural antibodies may facilitate clearance, the limited role observed for the C4-dependent classical pathway in this study suggests this defense mechanism may have limited efficacy, although it is possible adaptively-acquired antibodies may invoke this pathway.

The requirement for C4 and CD21/35 in controlling gastrointestinal colonization represents a novel and significant finding. While C4 deficiency is well-documented for its association with increased susceptibility to infections and autoimmune disorders—primarily due to its role in regulating humoral immunity^56–59^, the specific mechanisms by which complement components shape mucosal B cell responses remain poorly understood. Our findings in C4- and CD21/35-deficient mice reveal a strong and selective requirement for these components in limiting bacterial burden during antibiotic-induced supercolonization. This distinction underscores the hierarchical and context-dependent roles of the immune system in managing gut colonization, where the need to preserve a diverse commensal microbiota must be balanced with effective barrier surveillance and pathogen clearance. Given that *Kpn* is a leading source of hospital-acquired infections^60^, work of this nature is pivotal for devising strategies to limit colonization and dissemination events within susceptible patients and to mitigate transmission and outbreaks.

## Materials and Methods

### Mice

WT C57BL/6, C3⁻^/^⁻ (B6.129S4-C3tm1Crr/J), C4⁻^/^⁻ (B6.129S4-C4btm1Crr/J), and B6.129S7(NOD)-Cr2tm1Hmo/J mice were obtained from Jackson Laboratory. C4⁻^/^⁻ mice were further backcrossed two generations onto C57BL/6 mice. CD21/35-deficient mice were as previously described^41^. All mice were sex- and age-(8–10 weeks) matched, housed under sentineled SPF conditions, acclimated to BSL2 housing 24 hours prior to infections, and used under Wake Forest University IACUC approval. No exclusion criteria were applied.

### Infections

Oral infections were performed for sex- and age-matched cagemates of each genotype (n=4-5/group) at the same time to limit confounding factors, as described by Young et al.^30^. No randomization or blinding was applied. Mice were fasted for ∼4 h, then given ∼10⁶ CFU of streptomycin-resistant *Kpn* (KPPR1S^61,62^) in 2% sucrose-PBS, via two 50-µl doses one hour apart. For antibiotic-induced supercolonization, antibiotics were provided in drinking water three days prior to inoculation—streptomycin sulfate (500 mg/L) for KPPR1S infections and ampicillin (250 mg/L) for MKP103^36^. After 4 h fasting, mice received 10⁷ CFU of KPPR1S or MKP103 in 100 µl 2% sucrose-PBS via 20-gauge gavage. Feces and tissues were homogenized in PBS, centrifuged, and supernatants were serially diluted and plated on selective agar to assess bacterial burden.

### Cobra venom factor (CVF) treatment of wildtype mice

12.5 ug CVF per 25 grams of body weight were given to WT mice via intraperitoneal injection^21^ one day before infection and on days 1, 3, 5 and 7 after infection. Control mice received PBS injections.

### ELISAs

Fecal supernatants (100 mg/mL in PBS) were centrifuged at 14,000 × g for 10 min at 4°C and stored at –20°C. C3 and C4 levels were measured using ELISA kits (Abcam ab157711; Hycult HK217).

### *In vitro* complement killing and deposition assay

To assess complement-mediated killing, *Kpn* strains (KPPR1S and ΔwcaJ) grown in LB or cecal filtrate^63,64^ were pellet and resuspended to 4 × 10⁹ CFU/mL of PBS. Bacteria (∼2 × 10⁷ CFU; 5 µL), DMEM (45 µL) and either 5 µL of PBS, rabbit complement, or heat-inactivated complement were combined. After 30 min at room temperature, survival was assessed by serial dilution and plating on LB agar. To quantitate C3b deposition, KPPR1S was labeled with 1 µM CellTracker Violet (CTV, 30 min), washed in PBS + 1% BSA, and incubated with 10% rabbit complement, heat-inactivated complement, or PBS in RPMI at 37°C for 1 h. After washing, bacteria were stained with goat anti-C3/C3b F(ab’)2-FITC (25 min), washed, and fixed in 1% buffered formalin. Gating was based on size (Apogee beads) and CTV^+^ signal. Background staining was determined using goat IgG F(ab’)2-FITC (Supplemental Fig. 4).

### Phagocyte recruitment

Phagocyte recruitment was assessed 48h post-infection. After perfusion with 30 mL PBS, colons were excised, flushed, and digested in collagenase IV (1 mg/mL) and DNase I (5 U/mL) in PBS containing 2% newborn calf serum (NCBS) at 37 °C for 20–30 min with agitation, then mechanically disrupted and filtered through 70 μm mesh. Cells were pelleted (400 × g, 10 min), resuspended in PBS-2% NCBS containing Countbright beads, CD11b-BV650, CD45-AF700, GR1-AF647, and Live/Dead Aqua (25 min.). After washing and fixation (1.5% formalin), samples were analyzed on a FortessaX20 using FlowJo software.

## Statistical analysis

Statistical analyses were performed using GraphPad Prism v10.2.3, with graphing of individual data to assess assumptions where parametric tests were used. Two-group comparisons used the Mann-Whitney U test. Multiple groups were analyzed by Kruskal-Wallis with Dunn’s test, one-way ANOVA with Bonferroni correction, or two-way ANOVA with appropriate post hoc tests. Survival was analyzed using the Log-rank (Mantel-Cox) test.

## Data availability

All data generated and analyzed are included in this published article and its supplementary information file. Raw data generated for the current study are available from the corresponding authors on reasonable request or will be made publicly available in a public repository (Dataverse).

## Acknowledgments

This work was supported by the NIH/NIAID (R21AI178595). H.L.D. and A.EA-S. were supported by NIH training grant T32AI007401. Additional support was provided by the National Center for Advancing Translational Sciences (UL1TR001420) and the NCI (P30CA012197). We thank Emma Bennett and Noah Nutter for assistance with sample processing.

## Author Contributions

KMH: Conceptualization, funding acquisition, supervision, formal analysis, and writing-original draft and editing. MAZ: conceptualization, supervision, funding acquisition, writing - review & editing. JDV-B: data curation, formal analysis, writing-original draft, review & editing. JEJ-G: data curation, resources, NAN, HLD: data curation, AEA-S: data curation, writing - review & editing. AAH: data curation.

