## Supplemental Figures 1-4 for "Divergent roles for complement components C3 and C4 in controlling *Klebsiella pneumoniae* gut colonization and systemic dissemination"

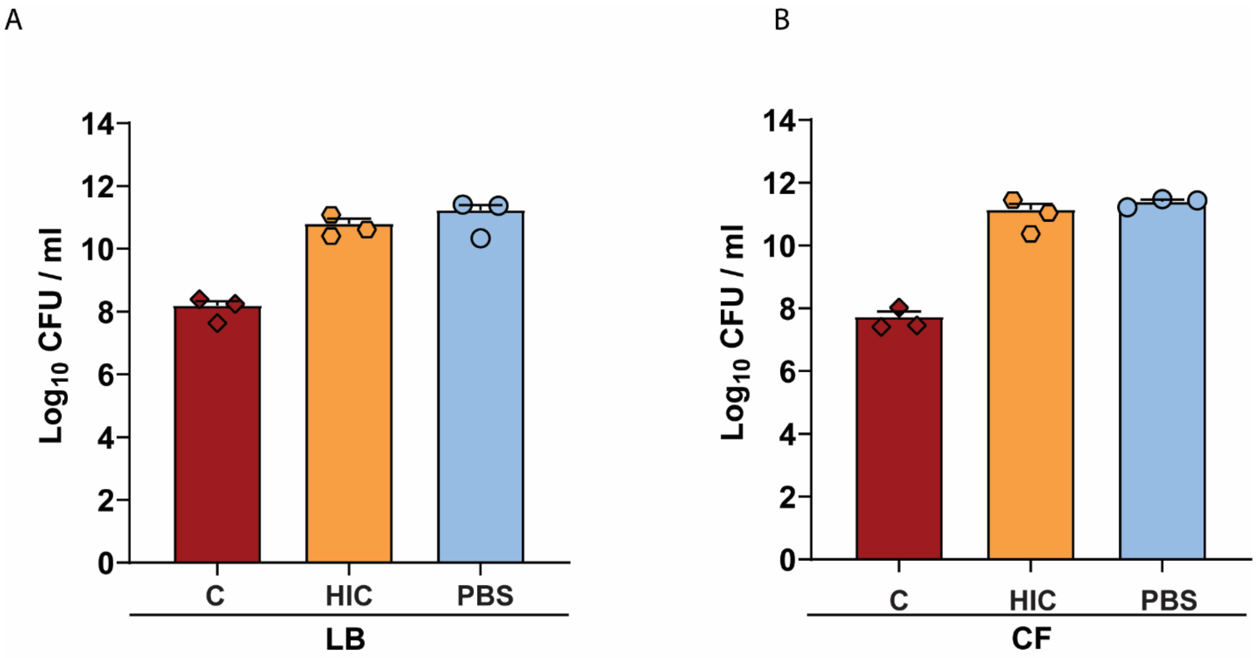


**Figure S1. Capsule protects *Kpn* against complement-mediated killing** (A–B) A *Kpn* isogenic capsule-deficient strain (*ΔwcaJ*) was incubated for 30 minutes at 37 °C in LB (A) or cecal filtrate (B) in presence of 10% rabbit complement, heat-inactivated complement, or PBS.

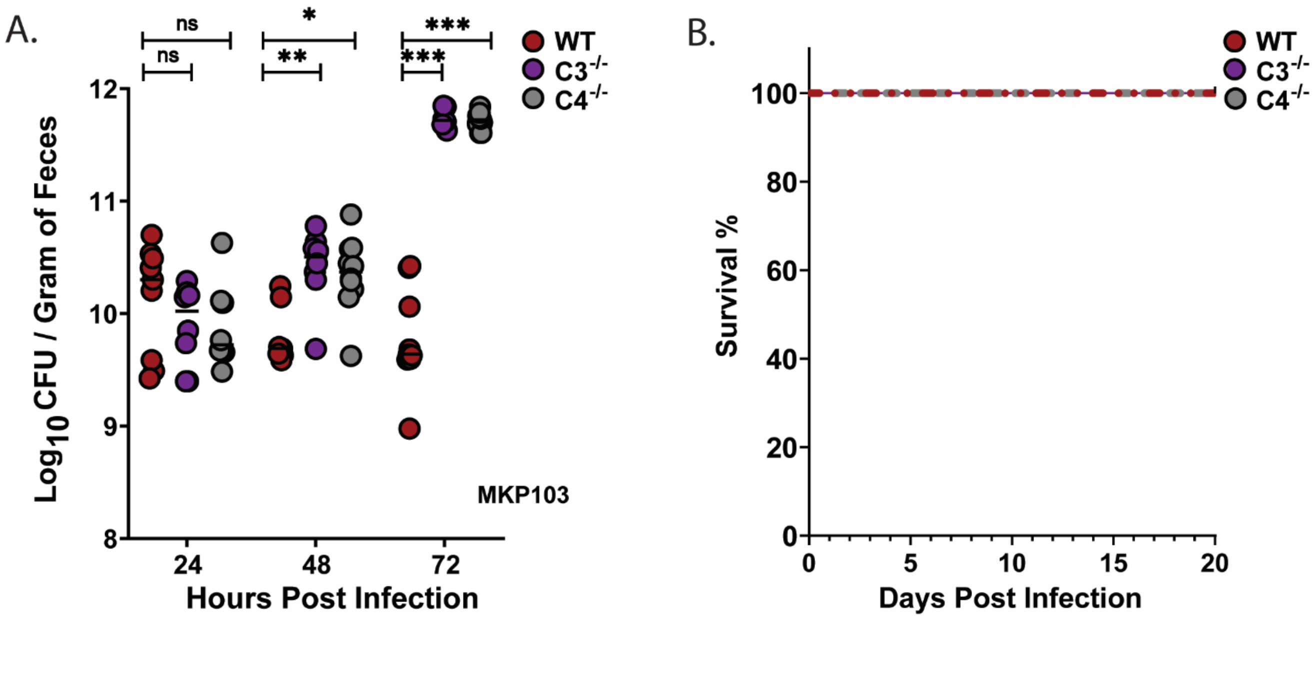


**Figure S2. Bacterial burden of Kpn classical strain following antibiotic pretreatment**. (A) Fecal shedding of mice colonized with MKP103 at 24-, 48-, and 72-hours post-gavage following ampicillin pretreatment (250 µg/mL in drinking water). Each data point represents an individual mouse; dotted lines denote the limit of detection (L.O.D.). All bacterial strains were grown overnight in LB, and mice were inoculated with ~$10^7$ CFU. Statistical significance was determined using the Kruskal–Wallis test with Dunn's post hoc test. n ≥ 9/group. *$P < 0.05$, **$P < 0.01$, ***$P < 0.001$, and ****$P < 0.0001$. (B) Kaplan–Meier survival curves of WT (red), $C3^{-/-}$ (purple), and $C4^{-/-}$ (gray) mice following MKP103 oral gavage.

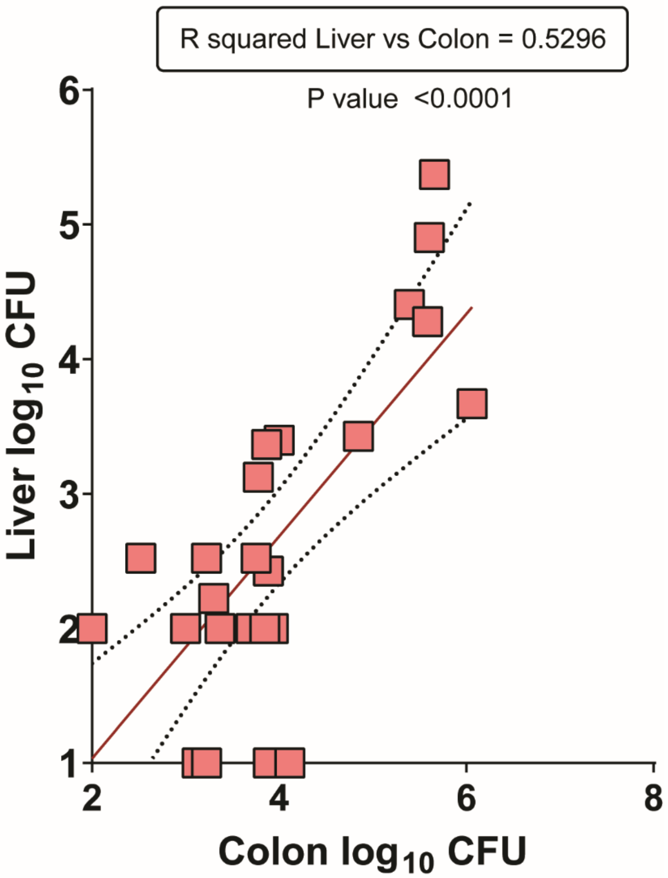


**Figure S3. Elevated bacterial burden in the gastrointestinal tract correlates with increased bacterial dissemination to extraintestinal sites in wild type mice.** (A) Correlation analyses of extra-intestinal organs (liver), bacterial burden of KPPR1S relative to gastrointestinal (colon) colonization at 24 hours PI in wild type mice, displayed as a simple linear regression curve. R squared for the lineal regression is listed, ($n \geq$ 27/group).

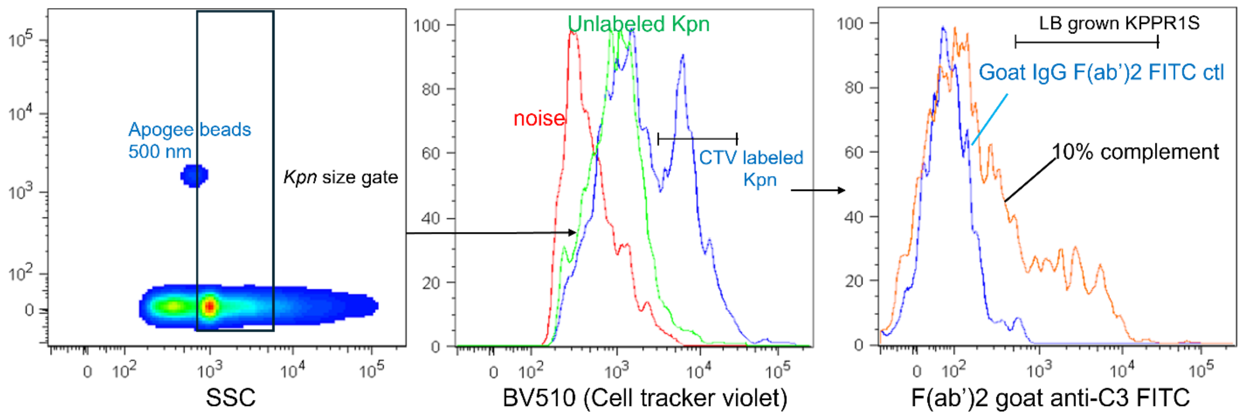


**Figure S4. Flow cytometry gating strategy to quantitate C3b deposition on KPPR1S.** *Kpn* was labeled with 1 uM Cell Tracker Violet for 30 minutes, washed with PBS containing 1% BSA, and incubated with 10% rabbit complement, 10% heat-killed complement, or PBS in RPMI at 37C for 1 hour. Bacteria was washed in PBS containing 1% BSA followed by staining with goat anti-C3/C3b F(ab')$_2$-FITC for 25 minutes. Bacteria were identified based on size using selective gating with Apogee beads followed by gating Cell tracker violet positive bacteria. Nonspecific goat IgG F(ab')$_2$-FITC was used to determine background staining. Mean fluorescent intensities (MFI) of C3b (FITC) signals were determined as a measure of C3b deposition level.
